# VRK1 is a Paralog Synthetic Lethal Target in VRK2-methylated Glioblastoma

**DOI:** 10.1101/2021.12.30.474571

**Authors:** Julie A. Shields, Samuel R. Meier, Madhavi Bandi, Maria Dam Ferdinez, Justin L. Engel, Erin E. Mulkearns-Hubert, Nicole Hajdari, Kelly Mitchell, Wenhai Zhang, Shan-chuan Zhao, Minjie Zhang, Robert Tjin Tham Sjin, Erik Wilker, Justin D. Lathia, Jannik N. Andersen, Yingnan Chen, Fang Li, Barbara Weber, Alan Huang, Natasha Emmanuel

## Abstract

Synthetic lethality — a genetic interaction that results in cell death when two genetic deficiencies co-occur but not when either deficiency occurs alone — can be co-opted for cancer therapeutics. A pair of paralog genes is among the most straightforward synthetic lethal interaction by virtue of their redundant functions. Here we demonstrate a paralog-based synthetic lethality by targeting Vaccinia-Related Kinase 1 (VRK1) in Vaccinia-Related Kinase 2 (VRK2)-methylated glioblastoma (GBM). VRK2 is silenced by promoter methylation in approximately two-thirds of GBM, an aggressive cancer with few available targeted therapies. Genetic knockdown of VRK1 in VRK2-null or VRK2-methylated cells results in decreased activity of the downstream substrate Barrier to Autointegration Factor (BAF), a regulator of post-mitotic nuclear envelope formation. VRK1 knockdown, and thus reduced BAF activity, causes nuclear lobulation, blebbing and micronucleation, which subsequently results in G2/M arrest and DNA damage. The VRK1-VRK2 synthetic lethal interaction is dependent on VRK1 kinase activity and is rescued by ectopic VRK2 expression. Knockdown of VRK1 leads to robust tumor growth inhibition in VRK2-methylated GBM xenografts. These results indicate that inhibiting VRK1 kinase activity could be a viable therapeutic strategy in VRK2-methylated GBM.

## INTRODUCTION

Recent years have witnessed a profound interest in targeting vulnerabilities in cancer stemming from synthetic lethal interactions – an approach to cancer treatment that specifically targets cancer cells while sparing normal healthy cells, thus increasing the therapeutic index of the therapeutic agent (1–3). The success of poly (ADP-ribose) polymerase 1 (PARP-1) inhibitors which are synthetic lethal with BRCA1 and BRCA2 mutations as well as other “BRCA-like” defects in homologous recombination demonstrated the potential of this therapeutic approach (4), and a subsequent largescale cancer dependency map (DepMap) resulted in discovery of additional novel synthetic lethal relationships (5–8). Using our proprietary bioinformatics pipeline, referred to as Tango Cancer Dependency Map (TANDEM), we analyzed public functional genomics data including the Cancer Cell Line Encyclopedia (CCLE) and identified one such novel paralog synthetic lethality, wherein VRK2-methylated GBM cell lines w19ere sensitive to loss of VRK1.

VRKs are a family of serine/threonine kinases that play a role in regulating transcription factors, chromatin remodeling, nuclear envelope formation, and cell cycle progression (9). There are three members of the VRK family – VRK1, VRK2 and a pseudokinase VRK3. VRK1 is found in both the nucleus and cytosol, and VRK2 localizes to the endomembrane of the endoplasmic reticulum and nuclear envelope (9). Functionally, VRK1 phosphorylates multiple substrates involved in both cell cycle progression and cell cycle arrest. Specifically, in response to mitogenic stimuli, VRK1 phosphorylates histones H3 and H2AX to facilitate chromatin remodeling, transcription factors ATF2, CREB, Sox2, and farnesoid X nuclear receptor HR1H4 to promote cell cycle progression, and BAF to regulate nuclear envelope formation (10,11). In response to stress signals, such as DNA damage, VRK1 phosphorylates p53, c-Jun, and 53BP1 to initiate cell cycle arrest for DNA damage repair (10,11). The functional role of VRK2 is less clear, however, it has been reported to downregulate apoptosis via direct interaction with anti-apoptotic protein Bcl-xL and by downregulating pro-apoptotic Bax (12). p53 and BAF have also been reported as substrates for VRK2 (13,14). Additionally, cells with low expression of VRK2 have shown enhanced sensitivity to chemotherapeutics (12).

The World Health Organization (WHO) classifies adult, diffuse gliomas into three types – astrocytoma, oligodendroglioma, and GBM (15). WHO further classifies gliomas based on isocitrate dehydrogenase (IDH) mutation status, thus allowing tumors to be classified considering both phenotype and genotype. GBM is the most common primary malignant brain tumor and is uniformly fatal due to minimal success with current and novel therapies (16). Approximately 90% of GBM is IDH-wildtype, typically primary GBM, and these patients have lower overall survival compared to IDH-mutant secondary GBM (17). O^6^-methylguanine-DNA methyltransferase (MGMT) promoter status is also used to stratify GBMs. Patients with methylated (or partially methylated) MGMT promoters are more likely to respond to the standard of care chemotherapeutic temozolomide, in contrast to those with unmethylated MGMT who are unlikely to benefit from the chemotherapeutic (18). In addition to temozolomide, the current standard of care includes surgical resection when possible and adjuvant radiation therapy, however, these treatments are associated with a median survival of only 15 months and a five-year survival rate of 6.8% (16,19,20).

Here we use CRISPR-based viability studies and cDNA rescue experiments to validate the paralog synthetic lethality between VRK1 and VRK2. Specifically, we show that in tumor cell lines with high VRK2 promoter methylation, and thus low VRK2 expression, knockdown of VRK1 induces cell death via G2/M arrest and DNA damage. We demonstrate that the kinase function of VRK1 is required for the synthetic lethality which posits that VRK1-targeting can be used as an approach to treat VRK2-methylated GBM.

## METHODS

### Cell Culture

Original cell lines were acquired from ATCC, ECACC, and JCRB. HAP1 isogenic cell lines were purchased from Horizon Discovery. All cell line stocks were routinely tested for mycoplasma. HAP1, LN229, YKG1, KNS60, U118MG, H4, LN18, T98G, YH13, KS1, KALS1, and SW1088 were maintained in Dulbecco’s modified Eagle’s medium (Gibco) supplemented with 10 % fetal bovine serum (FBS; GeminiBio) and 1 % sodium pyruvate (Gibco). U251MG cell lines were maintained in Eagle’s minimal essential medium (EMEM; Quality Biological Inc) supplemented with 10 % FBS. All cell lines were maintained in a cell culture incubator at 37 °C, 95% humidified air, and 5% CO_2_ atmosphere. For detailed information see supplemental methods.

### DNA Constructs and Cell Line Engineering

We used a dual vector lentiviral system for both CRISPR-Cas9 and tetracycline-inducible CRISPR-dCas9-KRAB cloning. All guide and cDNA sequences are reported in Supplemental Table 3. Lentivirus was generated by transiently transfecting Lenti-X 293T cells (Takara Bio) with lentiviral packaging mix (Cellecta), lipofectamine 3000 transfection reagent (ThermoFisher), and constructs diluted in Opti-MEM (Gibco). Virus was collected from the supernatant 48 hours post-transfection and filter sterilized using a 0.45 μm filter. Cells were infected with lentivirus and polybrene and selected in medium containing puromycin, blasticidin, or geneticin antibiotic as determined by the construct. For detailed information see supplemental methods.

### Colony Forming Assays

Cas9 expressing cells were seeded in tissue culture plates such that cells would reach 80-90% confluency in 14 days. The next day, cells were infected with lentivirus containing CRISPR guides and polybrene transfection reagent, and the following day, selected with 0.5 μg/mL puromycin. Cells were left to grow for 14 days and stained with crystal violet. For inducible CRISPR-dCas9-KRAB experiments, doxycycline was added to medium the day after seeding, and medium containing doxycycline was refreshed every 3-4 days during the 14-day growth period.

### Immunoblotting

Cells were rinsed in cold PBS and lysed in RIPA buffer (CST) supplemented with protease and phosphatase inhibitors (ThermoFisher) and universal nuclease (ThermoFisher). Lysates were cleared of insoluble material by centrifuging at 20,000 g for 10 min at 4°C and protein concentration was determined with the BCA Protein Assay (ThermoFisher). For immunoblotting 20-40 μg of protein in equal volumes were heated in LDS-sample buffer (Invitrogen) containing DTT for 5 min at 95°C. Samples were centrifuged at 20,000 g, separated by SDS-PAGE electrophoresis in 4-12% Bis-Tris gels (Invitrogen), and transferred to nitrocellulose membranes (Invitrogen). For detailed information see supplemental methods.

### Custom Antibody Generation

Phosphorylated BAF polyclonal antibody was raised in rabbits against the peptide N-MTTpSQKHRDFVAEPM by ProSci, Inc. (Poway, CA). Antibodies that recognized phosphorylated BAF was separated from total antibodies by a two-step immunoaffinity purification. Serum was first loaded on a column with immobilized non-phosphopeptide, flow through was then loaded onto a column with immobilized phosphopeptide, and finally purified antibody was eluted.

### Murine Xenograft Studies

The protocol was approved by the Institutional Animal Care and Use Committee (IACUC) of Pharmaron (Beijing, China) following the guidance of the Association for Assessment and Accreditation of Laboratory Animal Care (AAALAC). U251MG VRK2 low and U251MG VRK2 high cell lines were inoculated subcutaneously into 6- to 8-week-old female NOG mice (10 million cells / animal with 50% high density matrigel in EMEM) and allowed to form palpable tumors. Once the tumors reached ∼150 mm^3^, the mice were assigned to treatment groups with similar mean tumor volumes and treated with either saline or doxycycline (25 mg/kg) QD by oral gavage. Tumors were measured twice weekly throughout treatment.

### Cell Cycle Analysis

Cells were treated with doxycycline at the indicated concentrations and times, trypsinized, washed in PBS, fixed in 70% ethanol, and stained with Propidium Iodide/RNase Staining Buffer (BD Biosciences). Individual cells were characterized for forward and side scatter and DNA content was determined in 10,000 cells as measured by flow cytometry (FACS; excitation at 488 nm, emission measured using 600 nm bandpass filter) with an Attune cytometer (Thermo Fisher) or Novocyte cytometer (Agilent). Histograms and cell counts were generated using FlowJo X software.

### High-content immunofluorescence imaging

Cells were cultured in CellCarrier-96 Ultra microplates (Perkin Elmer) in the presence or absence of 1 μg/mL doxycycline for 5 days. Cells were fixed, blocked, and incubated with primary and secondary antibodies as indicated. Plates were imaged using Harmony high-content imaging and analysis software (Perkin Elmer). Briefly, nuclei were identified using the “find nuclei” function, and nuclei located at the periphery were removed using the “remove border objects” feature. Nuclear envelope roundness was quantified using Alexa-488 signal, and based on roundness, abnormal nuclear envelope positive cells were scored by the software. For detailed information, see supplemental methods.

### Phospho- and Total Proteomics Analysis

Cells were treated with and without doxycycline for the indicated times, washed twice in ice-cold PBS prior to harvesting. Proteins were digested using LysC/Trypsin and labeled for multiplexing using TMT labelling (ThermoFisher). Phospho proteins were collected using pSTY enrichment and fractionation and flow through from the pSTY enrichment was used for total protein-level analysis. All mass spectra were acquired on an Orbitrap Fusion Lumos coupled to an EASY nanoLC-1200 (ThermoFisher) liquid chromatography system. Heatmaps were generated on normalized scaled signal data using Morpheus Software (Broad Institute). Differential analysis between treatment groups was performed using an empirical Bayes method as in (21) and gene set enrichment analysis was calculated on proteins that were altered greater than +/- 2 Log fold-change. For detailed information, see supplemental methods.

## RESULTS

### VRK1 is a synthetic lethal target in VRK2-methylated cancer cell lines

To discover novel synthetic lethal interactions, we analyzed the large-scale cancer dependency database Achilles, where 808 cancer cell lines were screened with a genome-wide CRISPR Cas9 library to uncover genes essential for cell proliferation (22–24). We identified a subset of cell lines with low VRK2 expression that were sensitive to VRK1 knockdown (Fig 1A). Low VRK2 expression was primarily found in brain cancer and neuroblastoma cell lines, suggesting a neural lineage expression pattern. Genotype-Tissue Expression (GTEx) data reveals that normal human brain tissue has lower expression of VRK2 transcripts than other tissues, further suggesting a lineage-specific context (Fig S1A). Further analysis of CCLE data demonstrated that decreased expression of VRK2 strongly associated with increased promoter methylation (Fig 1B). VRK2 is rarely deleted or mutated in cancer cell lines or human tumors (Fig S1B-C) suggesting that the VRK2 methylation is the most likely cause of low VRK2 expression. To determine if VRK2-methylation occurs in primary human tumors, we analyzed data from The Cancer Genome Atlas (TCGA) low-grade glioma (LGG) and GBM datasets (25,26). Approximately two-thirds of LGG and GBM have low VRK2 expression that correlates with VRK2 promoter methylation, demonstrating that the “VRK2-low” genetic context is present in human brain cancer (Fig 1C). As noted above, current prognostic markers for LGG and GBM are MGMT promoter methylation status and IDH1/2 mutation status, so we queried whether VRK2-low status co-occurs with either of these markers. About 32% of LGG tumors with low VRK2 expression harbored mutations in IDH1 or IDH2, and approximately 55% of LGG and 21% of GBM tumors with low VRK2 expression had MGMT promoter methylation (Fig 1D, S1D).

**Figure 1.**
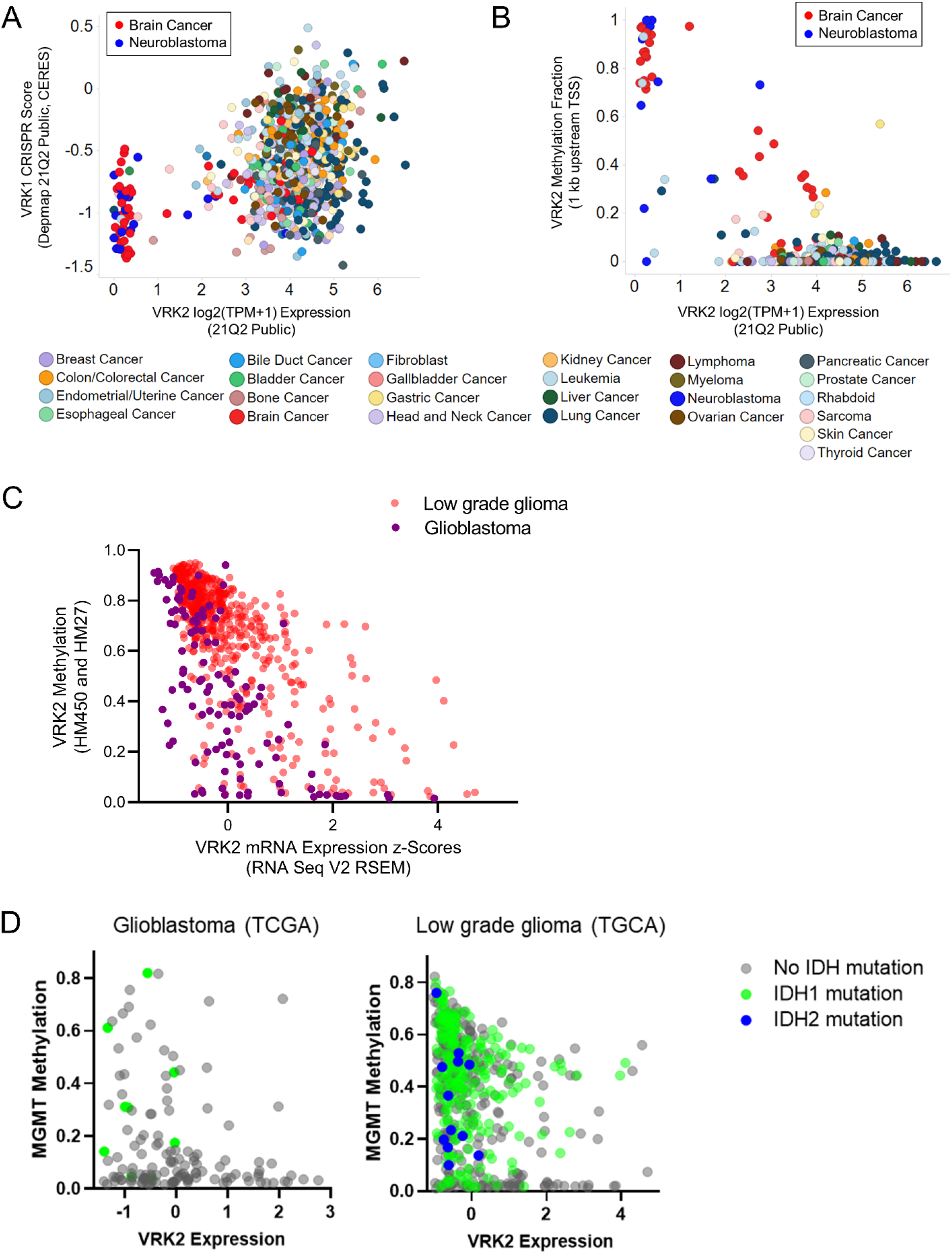
VRK2-methylated Glioblastoma and Neuroblastoma Cell Lines are Sensitive to VRK1 Loss. (A) Scatter plot depicting VRK2 expression and VRK1 CRISPR knockdown sensitivity score in 783 cancer cell lines. Color coded by primary lineage. (B) Scatter plot depicting VRK2 expression and VRK2 promoter methylation in 902 cancer cell lines. Color coded by primary lineage. (C) Scatter plot depicting VRK2 Expression and VRK2 Methylation for 530 low grade glioma (LGG; peach) and 116 glioblastoma (GBM; purple) tumors (D) Scatter plots with VRK2 expression and MGMT methylation for tumors as in (C), color coded by IDH mutation status.

### VRK1-VRK2 synthetic lethality *in vitro* and *in vivo* in GBM cell lines

To validate the synthetic lethal relationship between VRK1 and VRK2, we obtained a VRK2-wildtype and -null isogenic cell line pair derived from the HAP1 model. We generated Cas9 derivatives of these cell lines and knocked out VRK1 using three different sgRNAs. Knockout of VRK1 was lethal in the VRK2-null cell line in a 14-day colony forming assay, whereas the HAP1 VRK2 wildtype cells continued to proliferate (Fig 2A). To control for Cas9 efficiency, we used sgRNA to the pan-lethal gene PLK1 and observed cell death in both cell lines. Immunoblots for VRK1 and VRK2 demonstrated 60-80 % knockdown with all three VRK1 guides and confirmed VRK2 expression levels in the cell lines (Fig 2B). On-target knockdown was confirmed by rescuing the lethal phenotype in the HAP1 VRK2-null cells by expressing a CRISPR edit-resistant VRK1 cDNA (Fig 2C). To determine if the VRK1 kinase activity is important for the synthetic lethal interaction, we engineered VRK1-kinase dead and reduced activity mutations. Lysine 71 is the catalytic lysine in the VRK1 active site and the K71M mutation eliminated VRK1 kinase activity as determined by *in vitro* phosphorylation activity on a VRK1 substrate (Fig S2A). Edit-resistant VRK1 kinase dead (K71M) mutant and a previously published reduced activity mutant (K179E) (27) failed to rescue the anti-proliferative phenotype, indicating that the kinase activity of VRK1 is required for the synthetic lethal interaction. Re-expression of VRK2 in the VRK2-null cells also rescued the lethality confirming that VRK2 loss in the VRK2-null cell line is the cause of the anti-proliferative phenotype. Immunoblots for VRK1 and VRK2 confirm the expression of the respective cDNA constructs (Fig 2D). To further evaluate the synthetic lethal interaction in VRK2-methylated GBM cell lines, we tested the effects of VRK1 knockdown in a panel of VRK2-methylated and -unmethylated GBM cell lines (Fig S2B). VRK2-methylated cell lines were more sensitive to VRK1 knockdown than VRK2-unmethylated in a 14-day colony forming assay (Fig 2E-F). The data were quantified by normalizing colony intensity compared to intron cutting controls and were controlled for Cas9 efficiency by subtracting the colony intensity of PLK1 pan-lethal controls (Fig 2G). We further tested sensitivity to VRK1 CRISPR knockout in two non-GBM lines, RKO and SNU-398 and observed no proliferative defect in a similar colony forming assay (Fig S2C).

**Figure 2.**
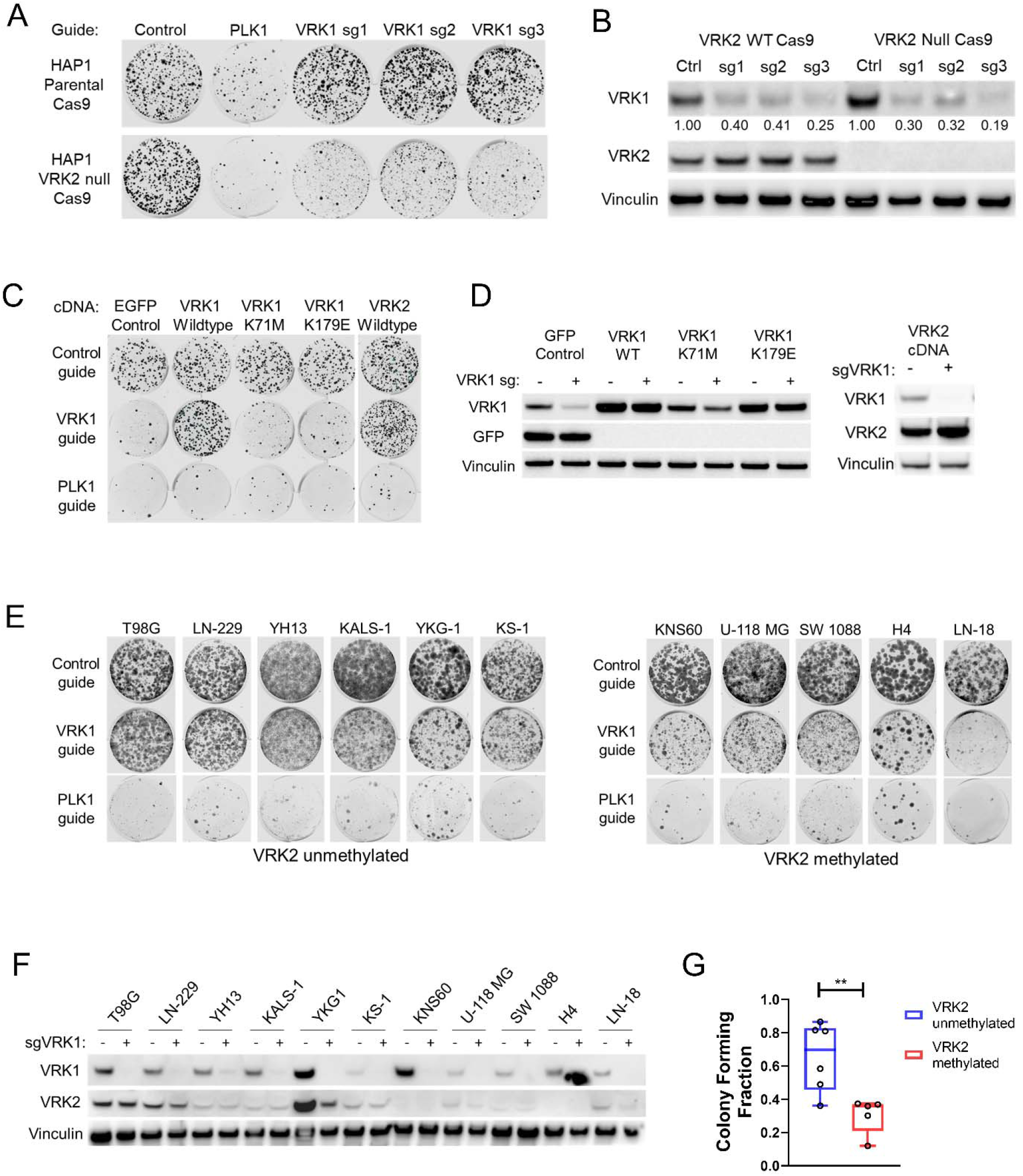
VRK1 is Synthetic Lethal in VRK2-null and VRK2 Methylated Cell Lines. (A) 14-day colony forming assay in HAP1 parental and VRK2-null cells with CRISPR knockdown for intron-cutting negative controls, positive control PLK1 and three VRK1 guides. (B) Immunoblots from (A) at 3 days. Quantification of VRK1 bands, normalized to Vinculin and relative to intron-cutting-controls, are indicated below the blot. (C) 14-day colony forming assays of VRK1 knockdown in HAP1 VRK2-null cell line with ectopic expression of the indicated cDNA constructs. (D) Immunoblots from (C) at 3 days. (E) 14-day colony forming assays of VRK1 knockdown in a panel of VRK2-unmethylated and -methylated GBM cell lines. (F) Immunoblots from (E) at 3 days depicting VRK1 and VRK2 protein expression. (G) Quantification of colony forming intensities of (E) from two biological replicates, corrected for Cas9 cutting efficiency, **p<0.005, two-tailed t-test

To extend our *in vitro* findings to the *in vivo* setting, we first established doxycycline-inducible VRK1 knockdown in the VRK2-methylated U251MG GBM model. Clonal U251MG cells expressing ubiquitous dCas9-KRAB and doxycycline-inducible CRISPR guides for the VRK1 promoter (henceforth referred to as “VRK2-low”) were sensitive to knockdown of VRK1 in a 14-day colony forming assay (Fig 3A). We generated “VRK2-high” derivatives of the clonal cell lines by re-expressing ubiquitous VRK2 cDNA and observed a full rescue of the anti-proliferative phenotype (Fig 3B). Protein levels of VRK1 and VRK2 were verified by immunoblotting (Fig 3C). Similarly, dox-inducible knockdown of VRK1 in a VRK2-unmethylated LN229 GBM cell line showed no proliferative defect *in vitro* (Fig S3A-B). To evaluate the synthetic lethal interaction *in vivo*, mice harboring established, subcutaneous xenografts of U251MG VRK2-low and VRK2-high derivative cell lines were treated with doxycycline or saline solution via oral gavage. A subset of tumors from all arms of the study were collected at seven days post-treatment and immunoblots demonstrated successful dox-induced VRK1 knockdown in both models and sustained VRK2 expression in the VRK2-high tumors (Fig 3D). Tumors in the VRK2-low derivative models regressed after 36 days on treatment compared to the vehicle-treated arm. Doxycycline treatment was stopped in five mice and continued in four mice at day 50 at which time the tumors started to regrow in both arms (Fig 3E). Immunoblots of these tumors at endpoint (day 60) reveal re-expression of VRK1 protein suggesting selection pressure to maintain VRK1 expression is required for survival *in vivo* in the absence of VRK2 expression (Fig 3G). Tumors in the VRK2-high group continued to grow in the presence of doxycycline compared to the saline arm (Fig 3F), suggesting that VRK2 is a key predictor of response to VRK1 inhibition in this model.

**Figure 3.**
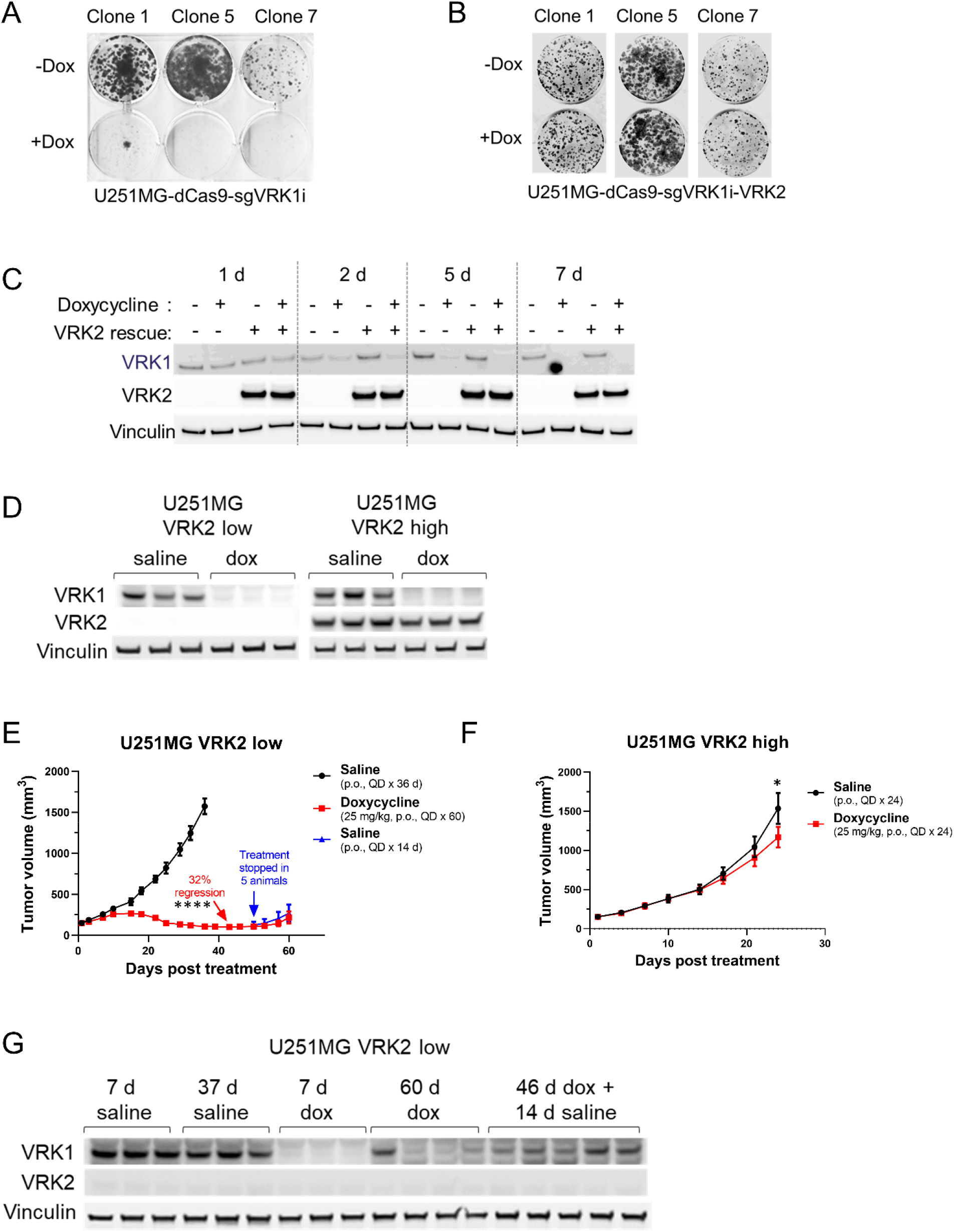
The VRK1-VRK2 Synthetic Lethality is Maintained *In Vivo*. (A) 14-day colony forming assays in U251MG VRK2-low cell line the absence or presence of 1 μg/mL doxycycline to induce VRK1 knockdown (B) Assay similar to (A) in the U251MG VRK2-high cell line (C) Immunoblots from (A) and (B) at 1-, 2-, 5- and 7-days post doxycycline treatment (D) Immunoblots from U251MG VRK2-low and VRK2-high xenografts in tumor-bearing mice treated with or without 25 mg/kg doxycycline QD for 7 days. (E) Tumor growth curves in mice bearing established 150 mm^3^ U251MG VRK2-low xenografts were treated with the indicated treatments. Data are presented as mean tumor volume ± SEM with 9 mice/data point up to Day 42, ****p < 0.0001 (two-tailed t - test). (F) Tumor growth curves as in (E) with U251MG VRK2-high xenografts. Data are presented as mean tumor volume ± SEM with 9 mice/data point, *p=0.032 (two-tailed t - test). (G) 7-day and endpoint immunoblots tumors from (E).

### VRK1 regulates BAF activity to maintain nuclear envelope integrity

Since VRK1 plays an important role in cell cycle progression (11), we queried whether VRK1 knockdown led to aberrant cell cycling in the VRK2-low context. Doxycycline-induced VRK1 knockdown for seven days provoked a G2/M arrest in the U251MG VRK2-low cell line as demonstrated by flow cytometry of propidium iodide-stained cells (Fig 4A, S4A). The VRK2-high U251MG derivative continued to cycle, suggesting that the mechanism of VRK1-VRK2 synthetic lethality involved a G2/M arrest. A similar G2/M arrest was also observed in the HAP1 VRK2-null, but not HAP1 VRK2-wildtype cells upon VRK1 knockdown (Fig 4B, S4B). Since VRK1 phosphorylates several substrates involved in mitosis, we profiled the phosphorylation status of the reported substrates histone H3 (T3, S10), p53 (T18) and BAF (S4). We were unable to detect any phosphorylation of p53 at Thr18 with several commercially available antibodies (data not shown). We did not observe significant alterations in the phosphorylation of Histone H3 at Thr3 and Ser10 (Fig S4C). As there is no commercially available antibody for the detection of phosphorylated BAF, we generated a custom polyclonal phospho-BAF (S4) antibody. Lambda phosphatase-treatment of HAP1 lysates demonstrates that this antibody recognizes a mixture of phospho-BAF and total-BAF proteins (Fig 4SD). Phosphorylation of BAF (S4) was depleted upon VRK1 knockdown and restored upon VRK2 re-expression (Fig 4C, S4C). A similar alteration in phosphorylation of BAF (S4) was observed in the HAP1 VRK2-null cell line with VRK1 knock-out (Fig 4D). Consistent with previously published data, total BAF was also depleted upon VRK1 knockdown in both cell line models (28,29). BAF, encoded by the *BANF1* gene, is a highly conserved chromatin binding factor that regulates post-mitotic nuclear envelope formation by linking nuclear envelope proteins to the DNA (30). Depletion of BAF results in aberrant nuclear envelope formation, nuclear blebbing, and multinucleation (31). Knockdown of VRK1 for five days led to abnormal nuclear envelope, nuclear lobulation, multinucleation and arrested mitotic spindles as determined by Lamin B immunofluorescent staining (Fig 4E). This phenotype was not due to a doxycycline effect since a non-targeting control guide did not produce nuclear envelope defects in the presence of doxycycline (Fig S4D). Re-expression of VRK2 rescued the abnormal nuclear envelope phenotype (Fig 4E-F) suggesting that the synthetic lethality depends, in part, on the VRK1 and VRK2 substrate BAF.

**Figure 4.**
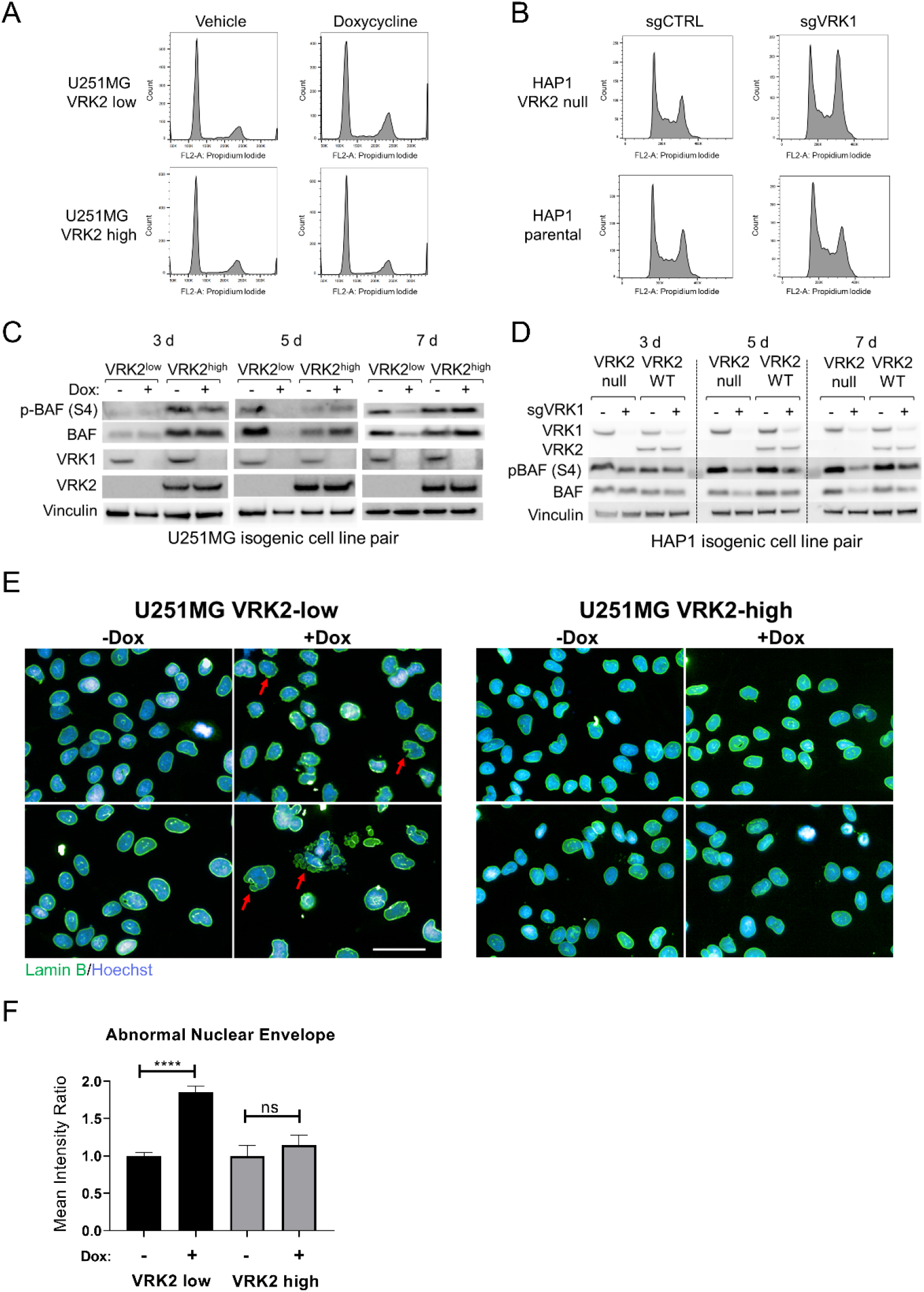
VRK1 Knockdown Results in G2/M Arrest, BAF Deregulation and Nuclear Lamina Defects. (A) Cell cycle distributions were determined by flow cytometric analyses in U251MG VRK2-low and VRK-high cells in the presence or absence of 1 μg/mL doxycycline for 7 days (B) Cell cycle distributions were determined by flow cytometric analyses in HAP1 VRK2-null and parental cells 5 days after VRK1 CRISPR knockdown (C) Immunoblots of U251MG VRK2-low and VRK2-high cells treated with or without doxycycline for 3-, 5- and 7-days. (D) Immunoblots of HAP1 VRK2-null and parental cells knocked out with VRK1 or intron-cutting control guides for 3-, 5-, and 7-days. (E) U251MG VRK2-low and VRK2-high cells were treated with or without doxycycline for 5 days and immunostained for Lamin B (green) and Hoechst (blue) and imaged by high-content imaging. Scale bar 50 μm. (F) Quantification of abnormal nuclear envelope from (E). ****p < 0.0001, ns= not significant (two-tailed t-test)

### The VRK1-VRK2 synthetic lethality activates the DNA damage response pathway

To further understand the mechanism underlying the synthetic lethality, we performed global phospho- and total-proteomic profiling in the U251MG VRK2-low and VRK2-high cell lines at five and seven days post-doxycycline treatment. Global changes in the proteome and phospho-proteome were more marked at seven days than five days, and thus, we performed differential analysis between VRK2-low and VRK2-high cell lines in the presence of doxycycline at the seven day timepoint in both datasets (Fig 5A, Supplemental Table 1). Consistent with our previous findings, differential analysis of proteomic profiling revealed notable GSEA enrichment of proteins involved in G2/M arrest such as PLK1, AURKA, AURKB, BIRC5, CDC6 and CCNB1 at seven days post doxycycline treatment (Fig 5B-C, Supplemental Table 2). The changes in G2/M proteins were observed as early as five days (Fig S5) and at the later seven day timepoint, we also observed an accumulation of proteins involved in DNA repair such as RAD51, PCNA, RPA3 and members of the RFC complex. The phospho-proteomics data also indicate a G2/M arrest due increased phosphorylation of CDK1 (CDC2) at Tyr15 which inhibits the progression of the cell cycle. We additionally observed increased phosphorylation of CDC20 at S41 and T70 which is reported to accumulate during mitosis (32). Consistent with the nuclear envelope defect in VRK2-low cells upon VRK1 knockdown, we observed an accumulation of phosphorylated Lamin A/C and TMPO which signal nuclear envelope breakdown (33). Phosphorylated BAF was not detected in this dataset likely due to the small molecular weight and low abundance of the protein. The G2/M and DNA damage response proteins were not altered in the VRK2-high cell line suggesting that these processes are involved in the VRK1-VRK2 synthetic lethal interaction. A subset of G2/M and DNA damage response markers were validated by immunoblotting (Fig 5E). These data demonstrate that the synthetic lethal mechanism of VRK1 perturbation in a VRK2-low GBM cell line is a G2/M cell cycle arrest and subsequent DNA damage.

**Figure 5.**
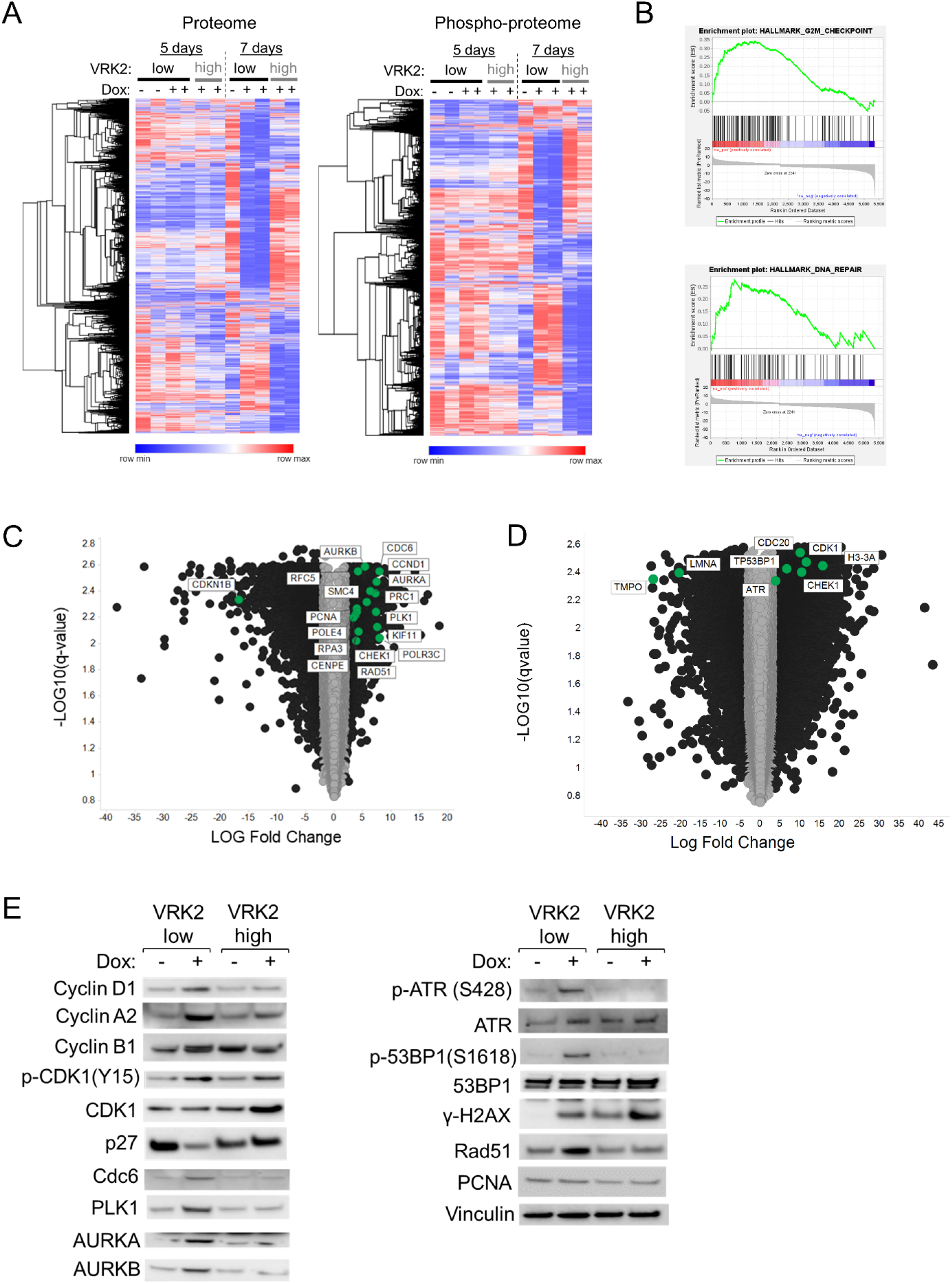
Phospho- and Total-Proteomics Reveals DNA Repair Pathways are Activated upon VRK1 Knockdown in VRK2-low Cells. (A) Heat maps showing total-proteomics (left) and phospho-proteomics (right) in VRK2-low and VRK2-high U251MG cells at 5- and 7-days post-doxycycline. (B) Gene Set Enrichment Analysis (GSEA) of 7-day total proteomics data in the doxycycline conditions. (C) Volcano plots differential expression analysis of total proteomic data. The x-axis represents log2 fold change and the y-axis represents the false discovery rate [-log10(q-value)]. Black circles – proteins with greater than +/- 2 log-fold change. (D) Volcano plots differential expression analysis of phospho-proteomic data. The x-axis represents log2 fold change and the y-axis represents the false discovery rate [-log10(q-value)]. Black circles – proteins with greater than +/- 2 log-fold change. (E) Immunoblots of select proteins from U251MG VRK2-low and VRK2-high cells treated with or without doxycycline for 7 days.

### Patient-derived GBM models maintain the VRK2-low state

Since available cell line models of GBM are not representative of the heterogeneous nature of the disease (34), we evaluated VRK2 expression levels was present in patient-derived GBM models. We performed immunoblotting for VRK2 in a panel of patient-derived GBM models and observed low VRK2-expression in seven out of nine models (Fig 6A). Measurement of VRK2 mRNA transcript levels by quantitative RT-PCR corroborated the immunoblot results (Fig S6). Since GBM contains self-renewing, tumorigenic cancer stem cells (CSCs) that contribute to tumor initiation and therapeutic resistance (35), we asked whether VRK2 expression differs between differentiating and non-differentiating media conditions. Each of the models were cultured either in CSC-(non-adherent, serum-free) or in non-CSC-(adherent, with serum) promoting conditions as previously described (36). VRK2 protein levels measured by immunoblotting remained unchanged in both media conditions suggesting stability of the promoter methylation regardless of differentiation status (Fig 6B). These data demonstrate that the VRK2-low context is maintained in patient-relevant GBM models.

**Figure 6.**
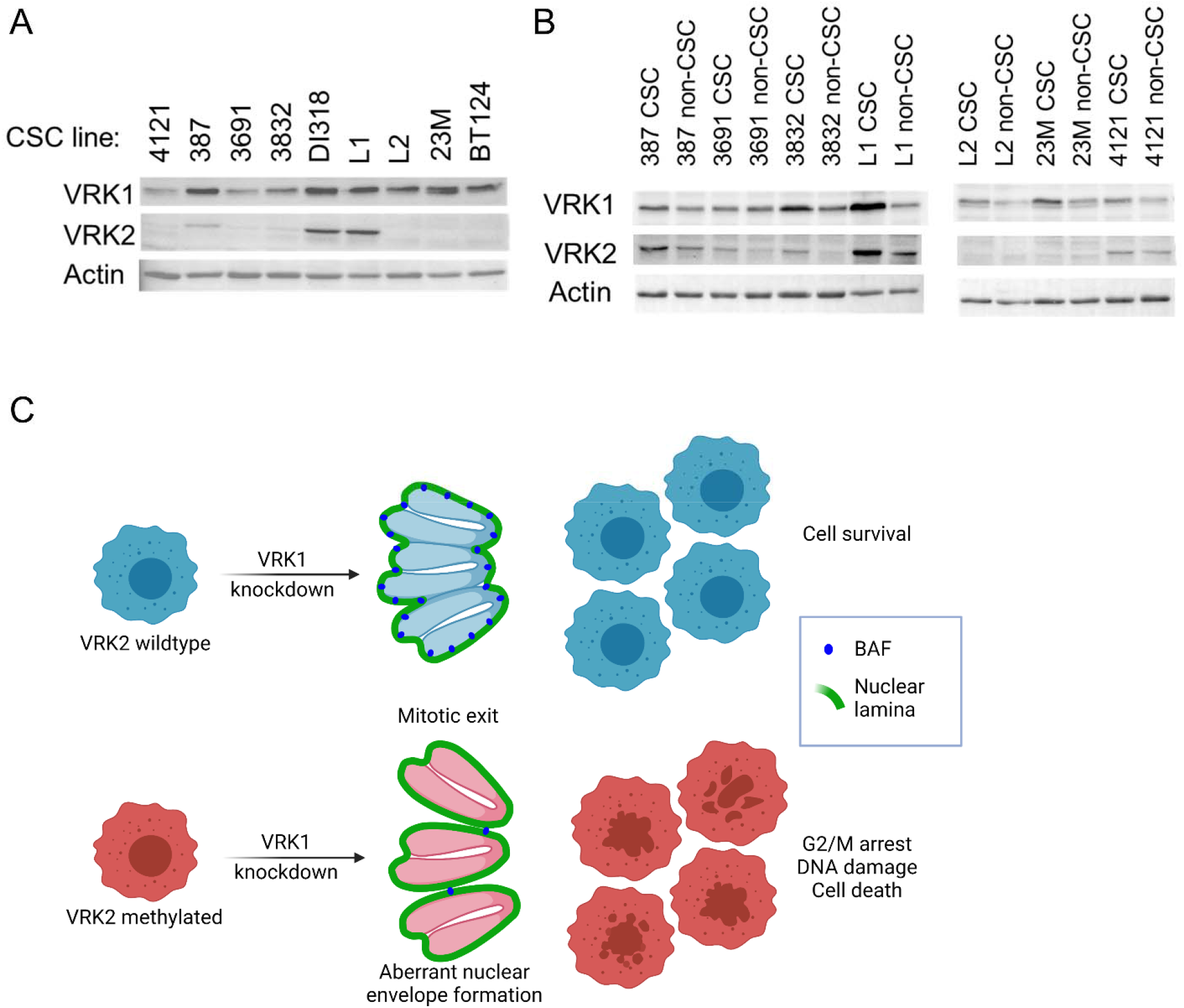
VRK2-methylated Context is Maintained in Patient-Relevant Glioblastoma Models. (A) Immunoblots from 9 GBM CSC models (B) Select CSC models were cultured in CSC or non-CSC medium and immunoblotted for VRK1 and VRK2 protein. (C) Schematic depicting mechanism of the VRK1-VRK2 synthetic lethality.

## DISCUSSION

Recent data from genome-scale CRISPR-Cas9 screening across hundreds of cancer cell lines have resulted in nomination of multiple novel targets for potential therapeutic development (37–40). In this study, we identified VRK1 as a paralog synthetic lethal target in VRK2-methylated GBM and neuroblastoma cell lines. Using CRISPR-based genetic tools, we demonstrate that knockdown of VRK1 in VRK2-null and VRK2-low expressing GBM cell lines is lethal, and results in defective nuclear envelope formation, G2/M arrest, and subsequent DNA damage. The synthetic lethal interaction is recapitulated *in vivo* in a VRK2-methylated U251MG GBM xenograft model, where VRK1 knockdown leads to tumor regressions. Xenografts from an isogenic VRK2-high U251MG cell line are insensitive to VRK1 knockdown, suggesting that the sensitivity depends solely on VRK2 expression. Importantly, our study demonstrates that the enzymatic activity of VRK1 is required for the VRK1-VRK2 synthetic lethality, which provides a path for small molecule drug discovery. Patient data indicates that the VRK2-methylated/VRK2-low context is present and may be common in LGG and GBM tumors. Taken together, these findings suggest that VRK1 is a promising synthetic lethal drug target in VRK2-methylated brain tumors, an aggressive indication with few therapeutic strategies currently available.

Past studies have identified synthetic lethal relationships among paralog genes such as SMARCA2-SMARCA4, ARID1A-ARID1B, and CREBBP-EP300 (41–43). The advantage of a synthetic-lethal therapeutic approach for cancer is the inherent large therapeutic index which maximizes anti-tumor efficacy while minimizing dose-limiting on-target toxicities (1). However, creating a paralog-selective inhibitor with these qualities hinges on developing a highly selective inhibitor that spares the non-target paralog despite nearly identical protein sequence homologies. Through mutant and wildtype cDNA rescue experiments, we show that the kinase function of VRK1 is essential in the synthetic lethal interaction. We believe that selectively targeting VRK1 over VRK2 may be possible as the kinase domains of VRK1 and VRK2 have approximately 80% protein sequence identity. Recent structural biology analysis revealed differential mechanisms for stabilization of an ATP-competitive inhibitor in the binding pocket of VRK1 compared to VRK2 (44,45), further supporting the possibility for development of a paralog-selective VRK1 kinase inhibitor.

VRK1 is one of the most abundant nuclear kinases in human cells and its overexpression is associated with poor prognosis in many solid tumors including GBM and neuroblastoma (46–49). We have shown for the first time that tumors with the genetic context of VRK2-methylation may benefit from selective inhibition or degradation of VRK1. Publicly-accessible cancer cell line data suggest that the VRK2-methylated context is restricted to cancer stemming from the neural lineage, i.e., GBM and neuroblastoma. Healthy human tissue expression data from the GTEx project further corroborates the lineage effect with lower VRK2 expression in neural-derived tissues compared to other tissues (Figure S1A). Additional evidence of a lineage effect comes from genetics on people with germline mutation of *VRK1*, which results in a neurological disease that manifests as prenatal microcephaly with pontocerebellar hypoplasia (50–53). Paralleling the human disease, mice with partial *Vrk1* knockdown have reduced brain weight and mild motor dysfunction (54). We posit the susceptibility of the developing brain to loss of VRK1 may be due to a naturally occurring synthetic lethal interaction stemming from reduced VRK2 expression in the neural cells compared to other lineages.

Although many substrates have been reported for VRK1 (55), our results demonstrate that BAF may be critical in the VRK1-VRK2 synthetic lethal interaction. We observe that knockdown of VRK1 leads to downregulation of BAF activity and results in a nuclear envelope defect that phenocopies BAF depletion (31,56–59). It is interesting to note that we and others observe decreases in both phosphorylated and total BAF, which may be due to protein instability in the absence of phosphorylation, a hypothesis that will require further testing (28,29). In addition to post-mitotic nuclear envelope assembly, BAF is involved in regulation of the DNA damage response and intrinsic immunity (60). Inherited germline *BANF1* mutations, which results in instability of the BAF protein, cause Nestor-Guillermo Progeria Syndrome, a premature aging disease characterized by genome instability and accumulation of DNA damage (61,62). Based on this study and previously published data, we postulate that the increased DNA damage response observed is due to genomic instability arising from the defective post-mitotic nuclear envelope. Interestingly, BAF also regulates cell-intrinsic immunity during mitosis by preventing cytosolic cGAS activation on genomic self-DNA (63). BAF dynamically outcompetes cGAS for DNA binding and prohibits the formation of DNA-cGAS complexes which are essential for inducing the type I interferon response. We did not observe cGAS activation or type I interferon response in our proteomics dataset suggesting that the pathway is not involved in the VRK1-VRK2 synthetic lethality. Taken together our study suggests that BAF plays a role in the VRK1-VRK2 synthetic lethal interaction. In VRK2-intact cells, VRK1 knockdown does not affect BAF activity due to compensation by VRK2, and results in formation of a smooth nuclear envelope at the end of mitosis. In VRK2-low cells, VRK1 knockdown leads to depletion of phosphorylated and total BAF, resulting in aberrant nuclear envelope with blebbing, lobulation and micronucleation. The cells arrest at the end of mitosis, accumulate DNA damage, and eventually die (Fig 6C).

Despite aggressive treatments for newly diagnosed GBM, almost all patients relapse with more aggressive tumors with minimal treatment options within one to two years. To date, efforts to develop treatments based on genetic alterations such as EGFR amplification, CDKN2A loss, TERT promoter mutation or PTEN mutation have been unsuccessful (64). Our findings indicate that VRK1 is a potential target for synthetic-lethal therapy in VRK2-low GBM, an aggressive indication with few therapeutic options. Interestingly, a recent study demonstrated that knockdown of VRK1 *in vitro* synergizes with temozolomide treatment by augmenting the DNA damage response (65). Together these data suggest that a VRK1 inhibitor may be used as a single agent or in combination with the current standard of care to augment therapeutic response.

The results of this study uncover a novel paralog synthetic-lethal interaction between VRK1 and VRK2 in GBM. We demonstrate that knockdown of VRK1 is lethal in VRK2-methylated GBM cell lines *in vitro* and *in vivo*, and the kinase activity of VRK1 is important in the interaction. The mechanism underpinning the lethality is BAF deregulation resulting in aberrant nuclear envelope formation, G2/M arrest, and subsequent DNA damage. These findings support the significant therapeutic potential of a VRK1 inhibitor in VRK2-methylated GBM.

## DATA AVAILABILITY

Mass spectrometry raw files will be uploaded to the UCSD MassIVE repository when accepted for publication.

## AUTHOR CONTRIBUTIONS

Conceptualization, F.L, A.H, and N.E.; Methodology, J.A.S., S.R.M, J.L.E., M.B., W.Z., M.Z., J.D.L., E.W., F.L., and N.E.; Formal Analysis, J.A.S., S.R.M, J.L.E., M.B., E.E.M-H., K.M., W.Z., F.L. and N.E.; Data Acquisition, J.A.S., J.L.E., M.B., E.M-H., K.M., W.Z., S.Z., F.L. and N.E.; Writing – Original Draft, J.A.S and N.E.; Writing – Review and Editing, N.E., J.L., J.N.A, and Y.Cs.; Visualization, N.E.; Supervision, J.L., E.W., J.N.A, Y.C., F.L, A.H., and N.E.

## ACKNOWLEDGEMENTS

We thank Dr. Douglas Whittington for structural analysis of VRK1 and Dr. Maria “Masha” Alimova for assistance with high-content imaging analysis.

---

**Supplemental Figure 1.**
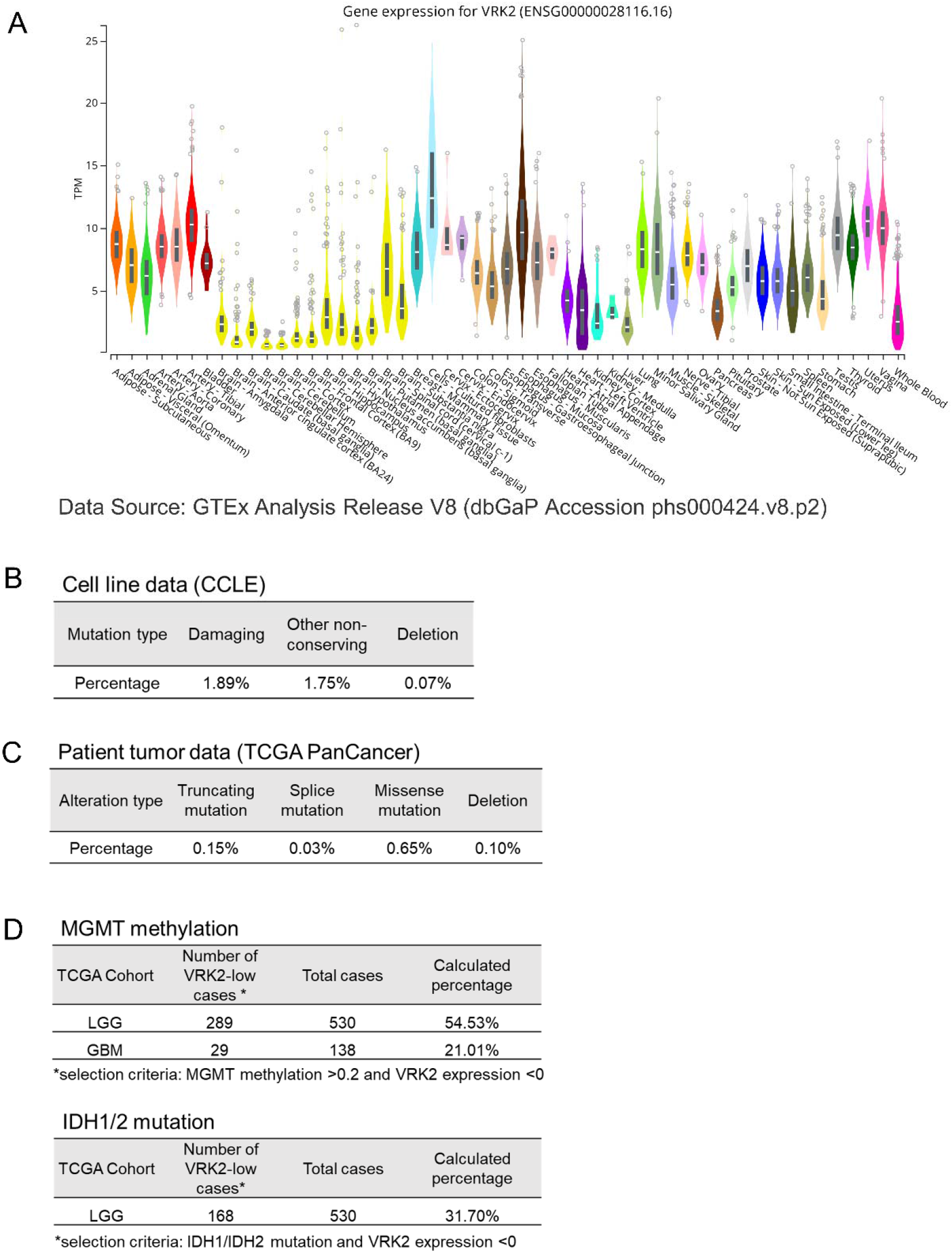
(A) VRK2 mRNA expression across all normal tissues (B) Percentage of VRK2 alterations in cancer cell lines (C) Percentage of VRK2 alterations in human tumors (D) Percentage of VRK2 low cases in MGMT-methylated and IDH mutant tumors

**Supplemental Figure 2.**
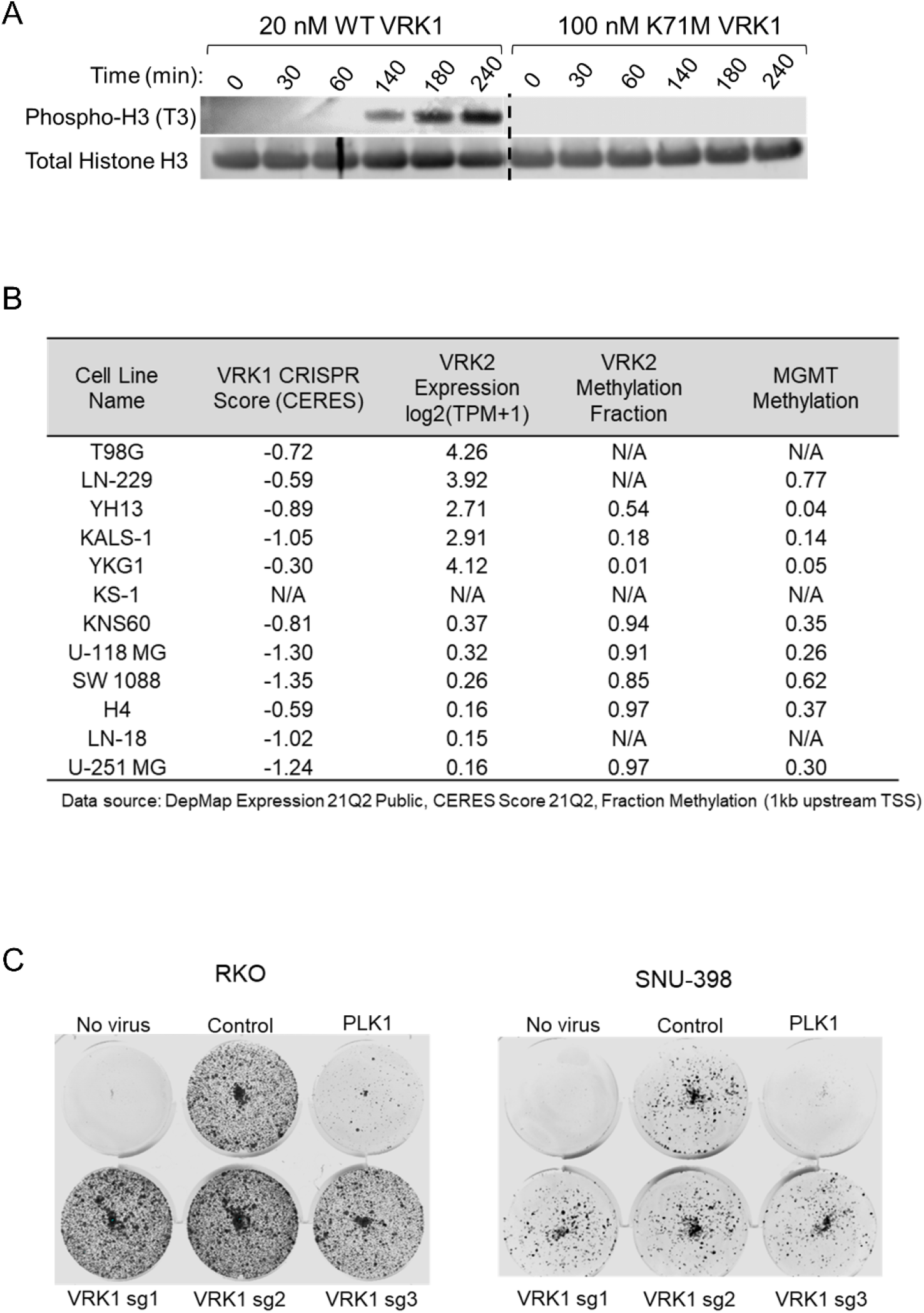
(A) Immunoblot depicting in vitro kinase activity of wildtype and mutant VRK1 on a histone H3 protein (B) Table showing CRISPR score, VRK2 expression, VRK2 methylation and MGMT methylation of a panel of GBM cell lines (C) 14-day colony forming assay as in Figure 2A of VRK1 CRISPR knockdown in two non-GBM cell lines.

**Supplemental Figure 3.**
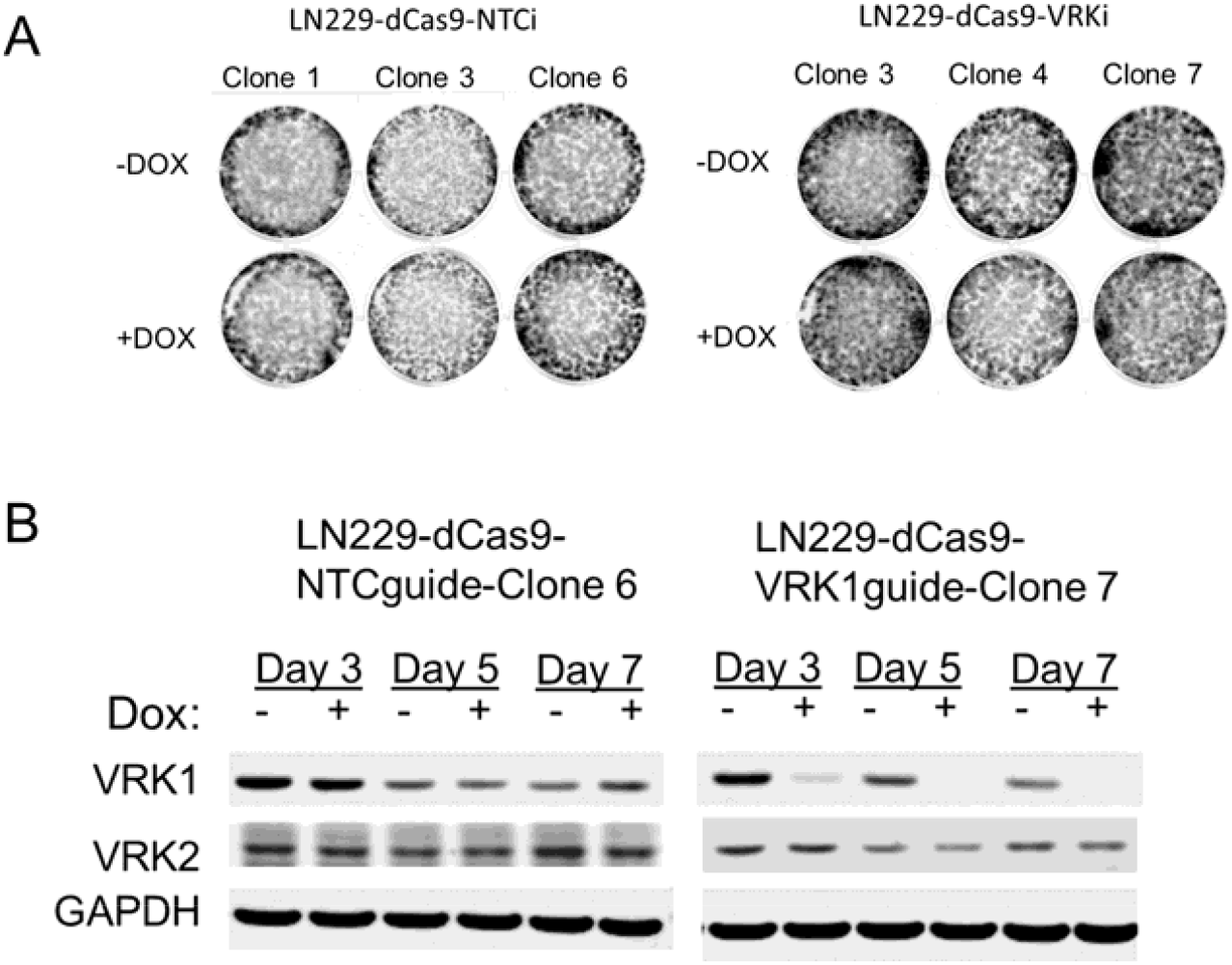
(A) 14-day colony forming assay of CRISPR-dCas9-KRAB inducible VRK1 knockdown (as in Figure 3A) of VRK2-high expressing LN229 clonal cell lines (B) Immunoblots of cells from (A) demonstrating VRK1 knockdown.

**Supplemental Figure 4.**
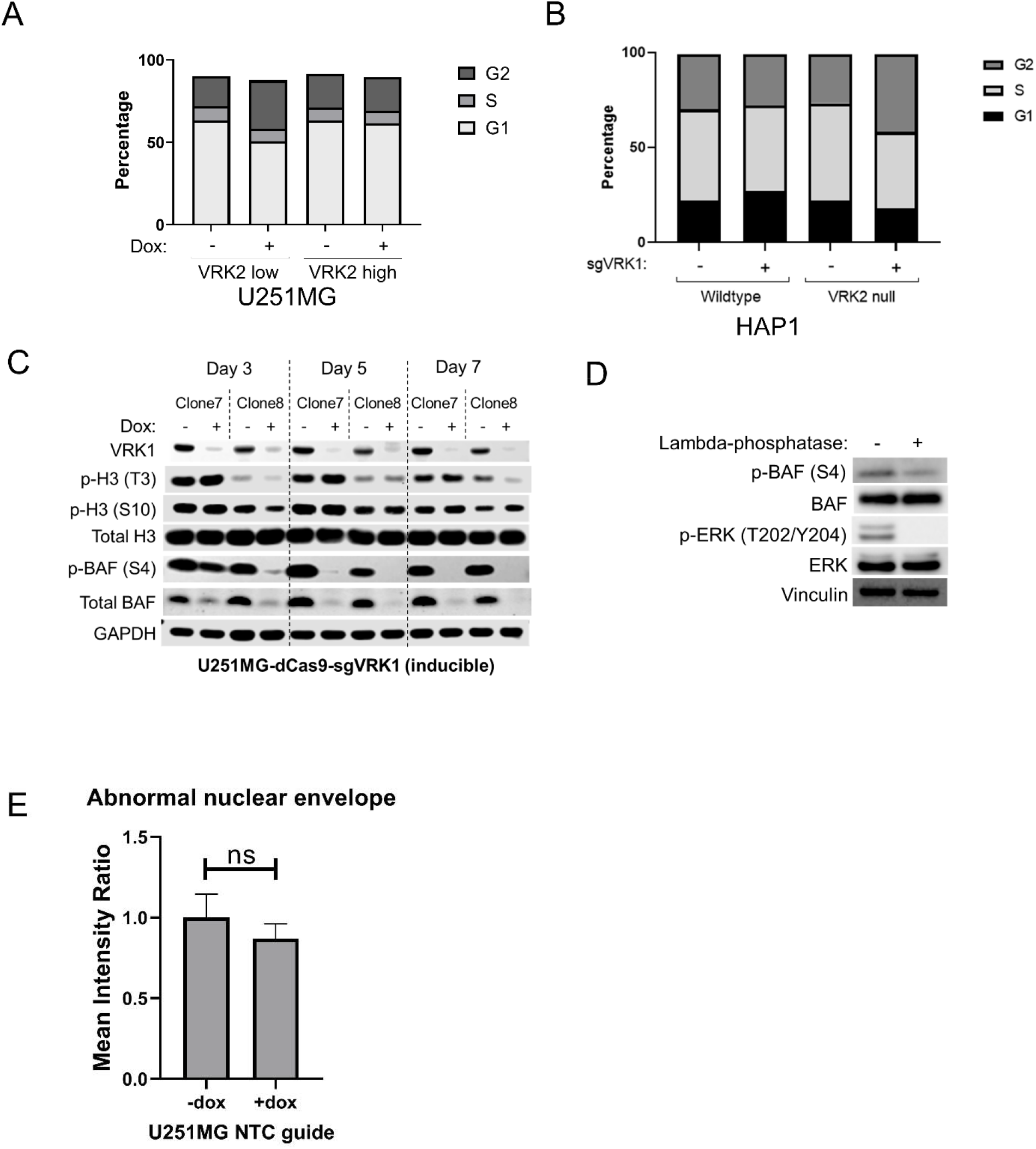
(A) Quantification of cell cycle distributions from Figure 4A (B) Quantification of cell cycle distributions from Figure 4B (C) Immunoblots of U251MG VRK2-low clonal cell lines at the indicated times (D) Immunoblots of HAP1 parental cell lysates treated with or without lambda phosphatase for 1 hour (E)Quantification of nuclear envelope phenotype in U251MG VRK2-low NTC control cell line treated with and without doxycycline.

**Supplemental Figure 5.**
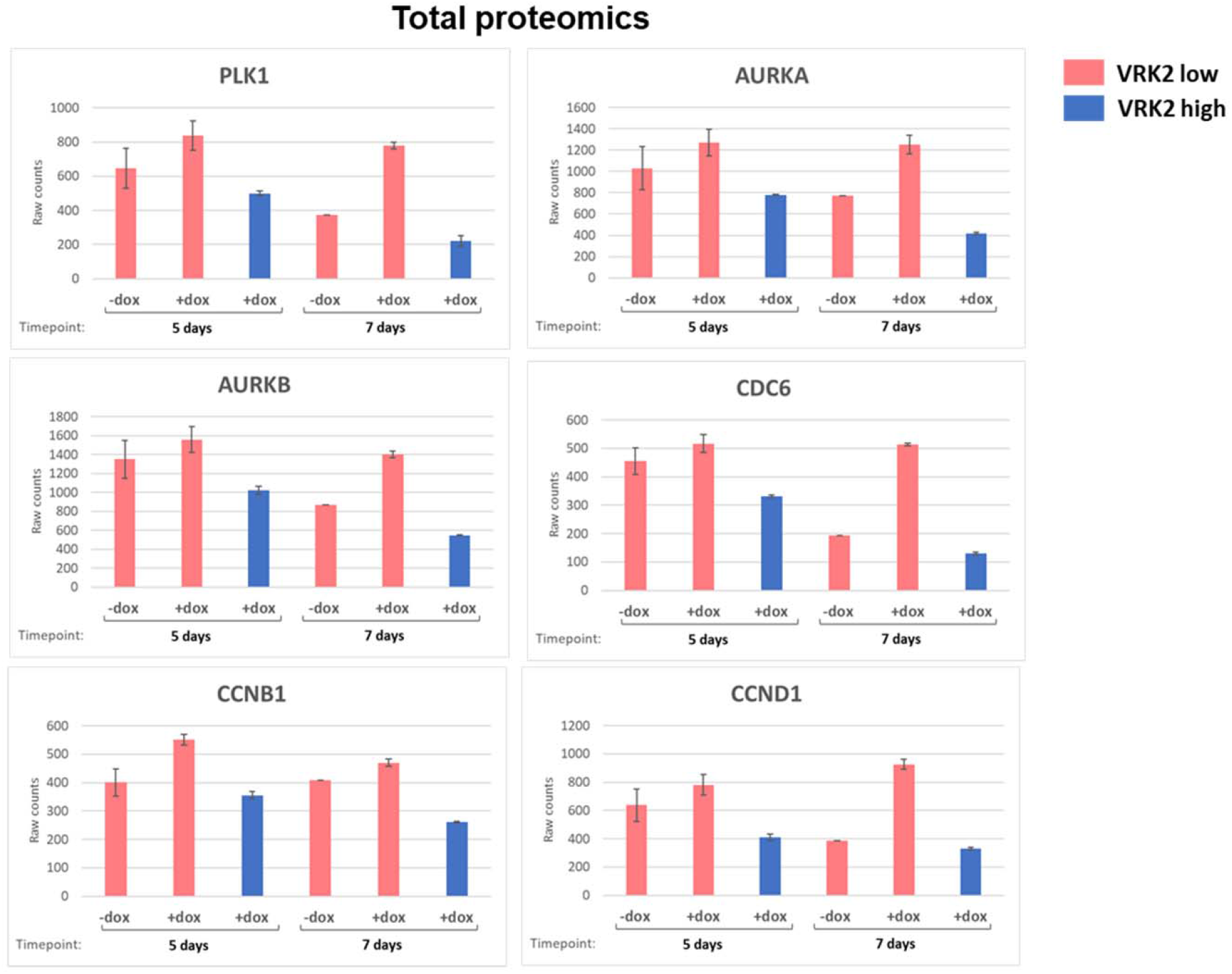
Total proteomics at 5- and 7-days post-doxycycline for cell cycle proteins.

**Supplemental Figure 6.**
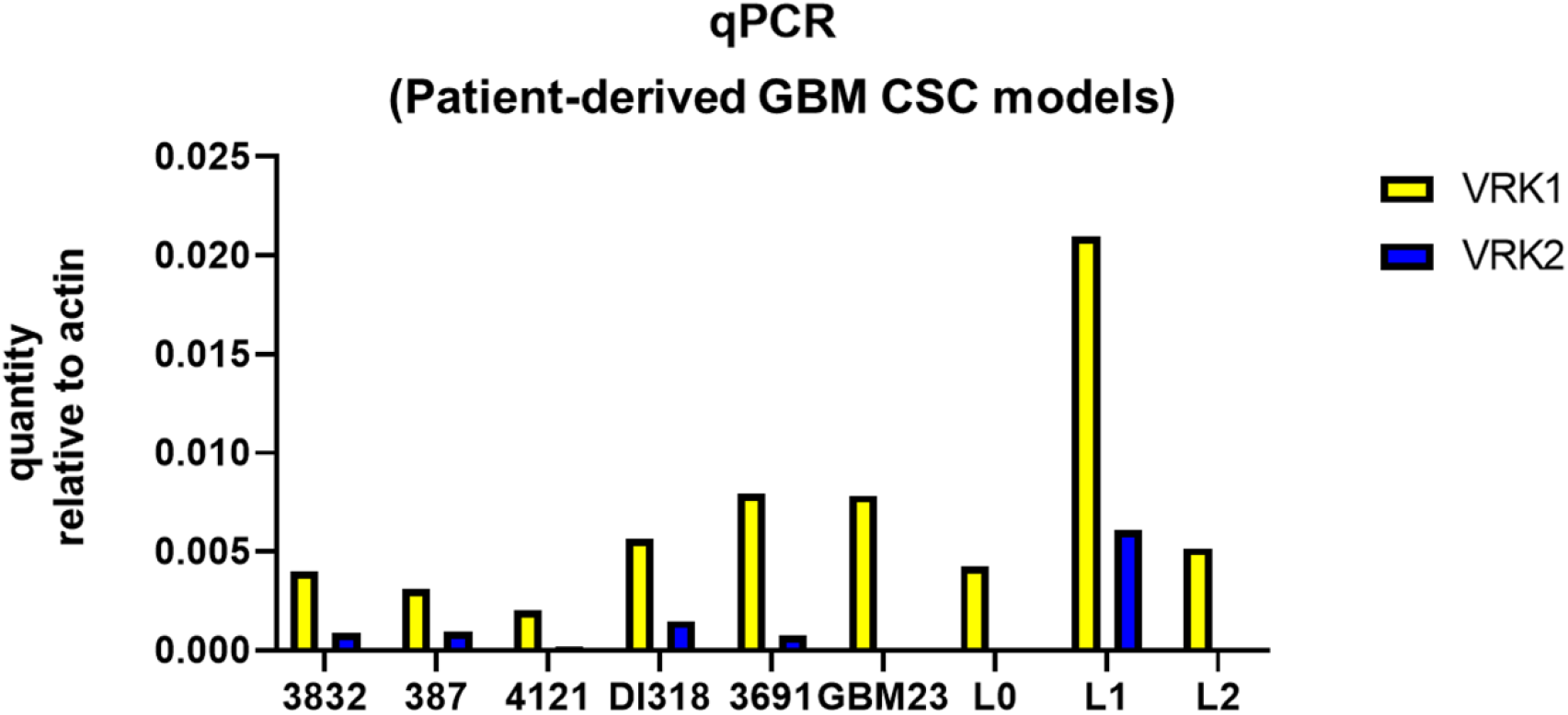
RT-qPCR for VRK1 and VRK2 from 9 CSC GBM models.

## Notes

### Competing Interest Statement

J.A.S., S.R.M., M.B., M.D.F., J.L.E., W.Z., S.Z., M.Z., R.T.T.S., A.H., B.W. J.N.A, E.W., Y.C., F.L. and N.E are employees and shareholders of Tango Therapeutics

