## Supplemental Methods for "VRK1 is a Paralog Synthetic Lethal Target in VRK2-methylated Glioblastoma"

**Cell Culture**

HAP1 isogenic cell lines were purchased from Horizon Discovery. Original parental cell lines were acquired from ATCC, ECACC and JCRB. All cell lines stocks were maintained in our in-house tissue culture bank and routinely tested for mycoplasma. HAP1, LN‑229, YKG‑1, KNS60, U‑118 MG, H4, LN‑18, T98G, YH13, KS‑1, KALS‑1 and SW 1088 were maintained in Dulbecco’s modified Eagle’s medium (DMEM; Gibco, 11965-084) supplemented with 10% fetal bovine serum (FBS; GeminiBio, 100-500) and 1X sodium pyruvate (Gibco, 11360-070). U-251 MG cell lines were maintained in Eagle’s minimal essential medium (EMEM; Quality Biological Inc, 112-018-101) media supplemented with 10% FBS. All cell lines were maintained in a cell culture incubator at 37°C, 95% humidified air, and 5% CO_2_ atmosphere. For all experiments using cell lines engineered with dCas9-KRAB, tetracycline-negative FBS (GeminiBio, 100-800) was used in place of regular FBS, and growth medium was supplemented with 1 µg/mL of doxycycline (Tocris, 4090) as needed. Growth medium for cell lines engineered with Cas9 was supplemented with 10µg/mL blasticidin (Gibco, 915438). Cas9 cell lines engineered with cDNA constructs also had medium supplemented with 1 mg/mL geneticin (Gibco, 10131‑027). Puromycin (Gibco, A11138‑03) at 0.5 µg/mL supplemented into growth medium to select for cells after introduction of guides.

**DNA Constructs and Cell Line Engineering**

We used a dual vector lentiviral system for both CRISPR-Cas9 and tetracycline‑inducible CRISPR‑dCas9‑KRAB cloning. All guide and cDNA sequences are reported in Supplemental Table 2. cDNA wildtype and mutant constructs were cloned using gBlocks purchased from Integrated DNA Technologies (IDT). All constructs were sequence verified using sanger sequencing. Lentivirus was generated by transiently transfecting Lenti-X 293T cells (Takara Bio) with lentiviral packaging mix (Cellecta, CPCP-K2A), lipofectamine 3000 transfection reagent (Thermo Scientific, L3000008), and cDNA/guide/Cas9 expression vectors in OPTI-MEM (Gibco, 31985-062). Virus was collected from the supernatant 48 hours post-transfection and filter sterilized using a 0.45 µm filter. Cells were infected with lentivirus and polybrene transfection reagent (Santa Cruz Biotechnology, sc‑134220) and selected in medium containing puromycin, blasticidin, or geneticin antibiotic as determined by the construct.

**Colony Forming Assays (CFAs)**

Cas9 expressing cells were seeded in tissue culture plates such that such that wells would reach 80-90% confluency 14 days post-selection. The next day, cells were infected with lentivirus containing CRISPR guides plus polybrene transfection reagent. The following day, infected cells were selected for using puromycin at 0.5 µg/mL. Cells were left to grow for 14 days post-selection, at which point they were stained using crystal violet. For inducible CRISPR‑dCas9‑KRAB experiments, doxycycline was added to medium the day after seeding, and medium containing doxycycline was refreshed every 3-4 days during the 14-day growth period.

**Immunoblotting**

Cells were rinsed in cold PBS (Gibco, 14190-136) and lysed in 1X RIPA lysis buffer (CST, 9806) supplemented with protease and phosphatase inhibitors (Thermo Scientific, 1861281) and universal nuclease (Thermo Scientific, 88700). Lysates were cleared of insoluble material by centrifuging at 20,000 g for 10 min at 4°C and protein concentration was determined with the BCA Protein Assay (Thermo Scientific, A53225). For immunoblotting 20-40 µg of protein in equal volumes were heated in LDS-sample buffer (Invitrogen, NP0007) containing DTT (CST, 7016) for 5 min at 95°C. Samples were centrifuged at 20,000 g, separated by SDS-PAGE electrophoresis in 4-12% Bis-Tris gels, and transferred to nitrocellulose membranes (Invitrogen, IB23001). Primary antibodies and dilutions used for immunoblotting are as follows: Actin (Bio-Rad, 12004163, 1:10000), BANF1 (Abcam, ab248281, 1:1000 and Abnova, H00008815-M01, 1:1000), GAPDH (CST, 2118S, 1:1000), GFP (CST, 2956S, 1:1000), Histone H3 (CST, 4499S, 1:1000), p-Histone H3 (S10) (CST, 53348S, 1:1000), p-Histone H3 (T3) (EMD Millipore, 07-424, 1:1000), p-BAF (S4) (ProSci, custom antibody, 1:500), Vinculin (Sigma, V9131, 1:3000), VRK1 (Santa Cruz Biotechnology, sc-390809, 1:500), VRK2 (Santa Cruz Biotechnology, sc-365199, 1:500). Secondary antibodies and dilutions used for immunoblotting are as follows: Anti-Mouse HRP (CST, 7076S, 1:2500), Anti-Rabbit HRP (CST, 7074S, 1:2500).

**Cell Cycle Analysis**

Cells were treated with doxycycline at the indicated concentrations and times, trypsinized, washed in PBS, fixed in 70% ethanol, and stained with Propidium Iodide/RNase Staining Buffer (BD Biosciences, 550825). Individual cells were characterized for forward and side scatter and DNA content was determined in 10,000 cells as measured by flow cytometry (FACS; excitation at 488 nm, emission measured using 600 nm bandpass filter) with an Attune cytometer (Thermo Fisher) or Novocyte cytometer (Agilent). In select experiments we used the Celigo (Nexelcom Bioscience) to analyze cell cycle. Briefly, cells were treated with doxycycline in 96-well plates, washed in PBS, fixed in 70% ethanol, and stained with Propidium Iodide/RNase Staining Buffer. Individual cells were characterized using the Celigo software for PI integrated intensity as a measure of DNA content. Histograms and cell counts were generated using FlowJo X software.

**High-content immunofluorescence imaging**

Cells were cultured at 500 cells per well in CellCarrier-96 Ultra microplates (Perkin Elmer) in the presence or absence of 1 µg/mL doxycycline for 5 or 7 days. Cells were fixed (4% PFA/0.25% Triton-X) for 15 minutes, washed three times with PBS and permeabilized (0.5% Triton-X in PBS) for 20 minutes. Cells were blocked (10% goat serum in PBS) for 1 hour at room temperature and incubated with primary antibody overnight at 4°C. Plates were washed three times with wash buffer (0.05% Tween-20 in PBS), incubated for 1 hour in secondary antibody and Hoechst 3342 (1:2000, Thermo, H3570) diluted in blocking buffer in the dark at room temperature, and washed three times with wash buffer and once with PBS prior to imaging. Primary antibody staining was performed with rabbit anti-LaminB1 (1:250, Abcam, ab16048) or rabbit anti-γ-H2AX (1:400, CST, 9718S). Secondary antibody used was fluorophore-conjugated anti-rabbit Alexa-488 (1:1000, Thermo, A11008).

Plates were imaged using Harmony high-content imaging and analysis software (Perkin Elmer). Briefly, nuclei were identified using the “find nuclei” function, and nuclei located at the periphery were removed using the “remove border objects” feature. Nuclear envelope roundness was quantified using Alexa-488 signal, and based on roundness, abnormal nuclear envelope positive‑cells were scored by the software.

**Phospho- and Total Proteomics Analysis**

All proteomics work was performed at IQ Proteomics (Cambridge, MA).

Cell lysis: Cell pellets were lysed in Dulbecco’s phosphate-buffered saline, pH 7.3 + 1% sodium dodecyl sulfate + HALT protease and phosphatase inhibitors (Thermo). Lysates were bead beat at 4°C using a Precellys Evolution homogenizer. Protein concentration in the lysates was quantified using the Pierce micro BCA assay (ThermoFisher Scientific). Proteins were reduced with 5 mM DTT and alkylated with 15 mM iodoacetamide. All protein from each sample was precipitated using methanol/chloroform. Following methanol/chloroform precipitation the protein pellet was washed three times with 100% methanol.

LysC/Trypsin Digestion and TMT labelling: The dried protein pellets were resuspended in 2 M guanidine-HCl, 100 mM EPPS, pH 8.0. Approximately 5 mg protein per sample was digested with LysC (Wako Chemicals USA) at a 1:25; protease:protein ratio and the proteins were digested at 25°C for 12 h. Following the LysC digestion, the guanidine-HCl concentration was diluted to 0.5 M with 100 mM EPPS, pH 8.0 and trypsin (Promega) was added at 1:50; protease:protein ratio. Trypsin digestion was carried out at 37°C for 8 h. Peptide concentrations were quantified using the Pierce peptide fluorescence assay (ThermoFisher Scientific). 3 mg of peptide per sample was labelled with TMT 11-plex reagent (ThermoFisher Scientific) at a 1.5:1 ratio (TMT reagent:peptide), incubating at 25°C for 3h. TMT reactions were quenched with 0.5% hydroxylamine and samples were acidified with TFA and combined into a “final mix”. The final mix peptides were desalted on 2 g Waters tC18 SepPak cartridges (Waters Corporation) and dried by centrifugal evaporation.

pSTY enrichment and fractionation: pY phosphopeptides were enriched using the Cell Signaling Technologies pY-1000 antibody kit as per the manufacturers protocol. pY enriched peptides were fractionated into 4 fractions (20%, 25%, 30% and 50% acetonitrile in 0.1% triethylamine) on Empore-C18 (3M) in-house packed StageTips prior to analysis by mass spectrometry. pY peptides were reconstituted in 5% formic acid + 5% acetonitrile for LC-MS3 analysis. The flow through from the pY enrichment was desalted on 2 g Waters tC18 SepPak cartridges (Waters Corporation) and dried by centrifugal evaporation. pST phosphopeptides were enriched using the Pierce Fe-NTA phospho-enrichment kit (ThermoFisher). In brief, the dried peptides from the pY flow through were resuspended in binding buffer provided with the kit. The peptides were bound and washed as per manufacturers protocol. Phosphopeptides were eluted from the Fe‑NTA resin with 50mM HK2PO4 pH 10.5. Labelled phosphopeptides were subjected to orthogonal basic-pH reverse phase fractionation on a 3x150 mm column packed with 1.9 µm Poroshell C18 material (Agilent) equilibrated with buffer A (5% acetonitrile in 10 mM ammonium bicarbonate, pH 8). Peptides were fractionated utilizing a 45 min linear gradient from 6% to 35% buffer B (90% acetonitrile in 10 mM ammonium bicarbonate, pH 8) at a flow rate of 0.8 mL/min. Ninety six fractions were consolidated into 24 samples and vacuum dried. The samples were resuspended in 0.1% TFA desalted on StageTips and vacuum dried. pST peptides were reconstituted in 5% formic acid + 5% acetonitrile for LC-MS3 analysis

Peptide fractionation for protein-level analysis: The flow through from the pST enrichment was dried by centrifugal evaporation. The dried peptides from the pST flow through were resuspended in 0.1% TFA. Approximately 125 µg of peptide mix was subjected to orthogonal basic-pH reverse phase fractionation on a 3x150 mm column packed with 1.9 µm Poroshell C18 material (Agilent) equilibrated with buffer A (5% acetonitrile in 10 mM ammonium bicarbonate, pH 8). Peptides were fractionated utilizing a 45 min linear gradient from 12% to 45% buffer B (90% acetonitrile in 10 mM ammonium bicarbonate, pH 8) at a flow rate of 0.8 mL/min. 96 fractions were consolidated into 24 samples and vacuum dried. The samples were resuspended in 0.1% TFA desalted on StageTips and vacuum dried. Peptides were reconstituted in 5% formic acid + 5% acetonitrile for LC-RTS-MS3 analysis.

Mass spectrometry analysis: All mass spectra were acquired on an Orbitrap Fusion Lumos coupled to an EASY nanoLC-1200 (ThermoFisher) liquid chromatography system. Approximately 2 µg of peptides were loaded on a 75 µm capillary column packed in-house with Sepax GP-C18 resin (1.8 µm, 150 Å, Sepax) to a final length of 35 cm. Peptides for total protein analysis were separated using a 90-minute linear gradient from 10% to 30% acetonitrile in 0.1% formic acid. The mass spectrometer was operated in a data dependent mode. The scan sequence began with FTMS1 spectra (resolution = 120,000; mass range of 350-1400 m/z; max injection time of 50 ms; AGC target of 5e5; dynamic exclusion for 60 seconds with a +/- 10 ppm window). The most intense precursors were selected within a 2 s cycle and fragmented via collisional-induced dissociation (CID) in the ion trap (normalized collision energy (NCE) = 35; max injection time = 35ms; isolation window of 0.7 Da; AGC target of 1e4). Following ITMS2 acquisition, a real-time search (RTS) algorithm was employed to score each peptide and only trigger synchronous-precursor-selection (SPS) MS3 quantitative spectra for high confidence scoring peptides as determined by a linear discriminant approach. Up to 8 MS2 product ions were further isolated and fragmented via high energy collisional-induced dissociation (HCD) with analysis in the Orbitrap (NCE = 55; resolution = 50,000; max injection time = 200 ms; AGC target of 7e4; isolation window at 1.2 Da for +2 m/z, 1.0 Da for +3 m/z or 0.8 Da for +4 to +6 m/z). pST peptides were separated using a 90-minute linear gradient from 5% to 23% acetonitrile in 0.1% formic acid. The mass spectrometer was operated in a data dependent mode. The scan sequence began with FTMS1 spectra (resolution = 120,000; mass range of 350-1400 m/z; max injection time of 50 ms; AGC target of 1e6; dynamic exclusion for 60 seconds with a +/- 10 ppm window). The ten most intense precursor ions were selected for ITMS2 analysis via collisional-induced dissociation (CID) in the ion trap (normalized collision energy (NCE) = 35; max injection time = 200ms; isolation window of 0.7 Da; AGC target of 2e4). Following MS2 acquisition, a synchronous-precursor-selection (SPS) MS3 method was enabled to select five MS2 product ions for high energy collisional-induced dissociation (HCD) with analysis in the Orbitrap (NCE = 55; resolution = 50,000; max injection time = 300 ms; AGC target of 1e5; isolation window at 1.2 Da for +2 m/z, 1.0 Da for +3 m/z or 0.8 Da for +4 to +6 m/z). pY peptides were separated using a 120-minute linear gradient from 7% to 26% acetonitrile in 0.1% formic acid. The mass spectrometer was operated in a data dependent mode. The scan sequence began with FTMS1 spectra (resolution = 120,000; mass range of 350-1400 m/z; max injection time of 50 ms; AGC target of 1e6; dynamic exclusion for 75 seconds with a +/- 10 ppm window). The ten most intense precursor ions were selected for FTMS2 analysis via collisional-induced dissociation (CID) in the ion trap (normalized collision energy (NCE) = 35; max injection time = 150ms; isolation window of 0.7 Da; AGC target of 3e4; m/z = 2-6; Orbitrap resolution = 15k). Following FTMS2 acquisition, a synchronous-precursor-selection (SPS) MS3 method was enabled to select five MS2 product ions for high energy collisional-induced dissociation (HCD) with analysis in the Orbitrap (NCE = 55; resolution = 50,000; max injection time = 300 ms; AGC target of 1e5; isolation window at 1.2 Da.

All mass spectra were converted to mzXML using a modified version of ReAdW.exe. MS/MS spectra were searched against a concatenated 2021 human Uniprot protein database containing common contaminants (forward + reverse sequences) using the SEQUEST algorithm (Eng et al., 1994). Database search criteria are as follows: fully tryptic with two missed cleavages; a precursor mass tolerance of 50 ppm and a fragment ion tolerance of 1 Da for peptide and phosphoserine and phosphothreonine (phosphotyrosine fragment ion tolerances were set to 0.02 Da); oxidation of methionine (15.9949 Da) or pSTY (79.9663304; pSTY searches only) was set as differential modifications. Static modifications were carboxyamidomethylation on cysteines (57.02146374) and TMT on lysines and N-termini of peptides (229.162932). Peptide-spectrum matches were filtered using linear discriminant analysis (Huttlin Cell 2010) and adjusted to a 1% peptide false discovery rate (FDR) (Elias Nat Methods 2007) and collapsed further to a final 1.0% protein-level FDR. Posttranslational modifications were localized using a probability-based algorithm similar to Ascore (Beausoleil SA, Nature Biotech. (2006) 24:1285-92). Proteins were quantified by summing the total reporter intensities across all matching PSMs.

**In Vitro Kinase Assay**

20 nM wild type VRK1 or 100 nM K71M mutant VRK1 were mixed with 2 µM recombinant Histone H3 protein (Active Motif, 31897) and 50 µM ATP (Promega, V703B) at ambient temperature to initiate the in vitro kinase assay. At each specified time point, the reactions were quenched by sample loading buffer (Invitrogen, NP0007). After running the protein gel and transferring to a nitrocellulose membrane, the proteins were blotted for using p-Histone H3 (T3) antibody (EMD Millipore, 05-746R, 1:500) or total Histone H3 antibody (Abcam, ab1791, 1:1000) with an overnight incubation. The membranes were then incubated with secondary antibody anti-rabbit Alexa 680 antibody (LI-COR, 926-68071, 1:5000) after wash. Detected signals were read and analyzed by LICOR Odyssey instrument.

**Statistical Analysis**

GraphPad Prism software was used for all statistical analysis. Where applicable two-tailed t-tests were used to determine statistical significance.

**In Vivo Studies**

The protocol was approved by the Institutional Animal Care and Use Committee (IACUC) of Pharmaron following the guidance of the Association for Assessment and Accreditation of Laboratory Animal Care (AAALAC). U-251 MG VRK2 low and U-251 MG VRK2 high cell lines were inoculated subcutaneously into 6- to 8-week-old female NOG mice (1×10^7^ cells / animal with 50% high density matrigel in EMEM) and allowed to form palpable tumors. Once the tumors reached ∼100 mm^3^, the mice were assigned to treatment groups with similar mean tumor volumes and treated with either saline or doxycycline (25 mg/kg) QD by oral gavage. Tumors were measured twice per week throughout treatment.
